# Prevalence and Molecular Detection of *Pasteurella multocida, Mannheimia hemolytica*, and *Bibersteinia trehalosi* in Sheep, Western Oromia, Ethiopia

**DOI:** 10.64898/2026.02.16.706086

**Authors:** Kenea Megersa Gemechu, Hambisa Abdi Bedassa, Demiso Merga Sima

## Abstract

Ethiopia has the biggest population of livestock in Africa, and small ruminants play a significant role in both meat consumption and revenue from the export of live animals and their skins. Infectious diseases, especially pneumonic pasteurellosis, are a key constraint in productivity, which is low despite their economic relevance for a variety of technological and non-technical reasons. Sheep are particularly susceptible to high rates of morbidity and mortality from the disease while under stress. These bacterial species were to be isolated, identified, and molecularly detected from sheep in certain regions of western Oromia, Ethiopia, that appeared to be healthy, as well as those that were clinically ill. Using multi-stage sampling, a cross-sectional study was carried out in three zones: Horro Guduru Wollega, East Wollega, and West Shawa, between January and December 2022. 384 sheep (220 healthy, 164 ill) had their nasal swabs taken, and they were analyzed bacteriologically, biochemically, and molecularly using PCR for the PHSSA, Rpt2, and CapA genes. To evaluate the relationships between risk factors and bacterial prevalence, data were examined using logistic regression and descriptive statistics. With P. multocida being the most commonly isolated species, followed by M. haemolytica and B. trehalosi, the overall prevalence of Pasteurellaceae was 21.1%. Pneumonic and young sheep had a greater prevalence, and there were significant correlations with age (OR = 2.41; 95% CI: 1.42–4.18) and illness status (OR = 3.47; 95% CI: 2.06–5.97), but not with sex or location. These results demonstrate the persistent risk of pneumonic pasteurellosis in Ethiopian sheep and the significance of species-level identification in directing focused interventions, such as management, treatment, and immunization plans. To molecularly describe isolates from other places and elucidate the pathogenic function of Pasteurella and Mannheimia species in illness development, more research is advised.

## Introduction

Ethiopia has the largest national livestock population in Africa, and livestock provides up to 40 % of agricultural gross domestic product, accounting for nearly 20 % of total GDP, and 20 % of national foreign exchange earnings ^1, 2^. Small ruminants have a significant role in Ethiopia’s livestock economy by providing over than 30% of all domestic meat consumption and generating income from exports of live animals, meat, and skin ^3^.

However, due to many technical and non-technical factors, productivity is marginal in country ^4^. Infectious diseases are among the technical factors, and from this, pneumonic pasteurellosis is the most common one ^5^.

There are two forms of pasteurellosis in small ruminants, namely pneumonic and septicemic pasteurellosis ^6^. *P. multocida, M. haemolytica*, and *B. trehalosi* are the most common causative agents of the disease, which cause 30% of deaths in feedlot cattle and acute outbreaks in sheep populations, resulting in huge mortality all across the globe ^7, 8, 9^. It is one of the leading causes of death in all age groups of sheep and is most commonly associated with stress ^10, 11^. The economic loss caused by ruminant pneumonia is estimated to be 8% of total production costs, which includes medical expenses, reduced food conversion, increased production costs, and reduced food supply for people ^12^.

Since Pasteurella, especially *P. multocida*, is part of the natural flora of the buccal-pharyngeal region, animals that are under stress, such as those transported, have respiratory infections, inclement weather, are poorly fed and ventilated, and are kept in crowded places, develop bacterial growth and proliferation, which later extend to the lower respiratory tract and cause pneumatic pasteurellosis ^13^.

Several studies on the species and its serotypes that cause pasteurellosis in sheep have been conducted in different countries of the world and different regions of Ethiopia, indicating that pasteurellosis is a major threat to sheep production.

Despite annual vaccination against pneumonic pasteurellosis with a monovalent vaccine (inactivated *P. multocida* biotype A produced at NVI), there is a high mortality and morbidity rate after respiratory distress in country ^14^. Therefore, due to the significant economic losses caused by mortality, morbidity, delayed marketing, unthriftiness among survivors, and high treatment costs, pasteurellosis is a high-priority issue at the national level. The main cause is the lack of a comprehensive study on the epidemiology of this disease, as well as the lack of cost-effective prevention and control methods best suited to different phenotypes and serotypes of the pathogen ^4^.

Even though sheep pasteurellosis is the main threat to sheep production across Ethiopia, and controlling pneumonic pasteurellosis is a difficult task that requires an understanding of its epidemiology, which in turn needs the ability to catalog the strains of the agents circulating in the country, there is no published scientific evidence indicating the species of Pasteurella that are common in Western Oromia. Therefore, this study aims to isolate and identify the species of Pasteurella from nasal swabs sampled from sheep in selected areas of western Oromia, Ethiopia.

## Materials And Methods

### Description of the Study Areas

The study was conducted in three selected western Oromia zones; these are Horro Guduru Wollega, East Wollega, and West Shawa zones of the Oromia regional state, Ethiopia. Oromia has a livestock population of 25,506,409 cattle, 9,752,385 sheep, 8,425,727goats, 1,309,923 horses, 120,161 mules, 3,900,800 donkeys, and 19,160,388 chickens ^15^.

### Study Animals

The study animals were local breeds of sheep. The study parameters were categorized into study sites (Horro Guduru Wallaga Zone, East Wallaga Zone, and West Shoa Zone), sexes (male and female), ages (young for those of one year or less, and adult for those of greater than one year, according to ^24^), and healthy status; diseased (sheep with signs of respiratory distress including an erratic breathing pattern, grunting on expiration, coughing, dyspnea, loss of appetite, lethargy, and serous to muco-purulent nasal discharges with fever) and apparently healthy sheep).

### Study Design and Sampling Method

A cross-sectional study design and multi-stage sampling methods were conducted to attain the objectives of the study from January 2022 to December 2022. Three zones were purposefully selected from western Oromia depending on the security of the area. In the second stage, from each zone, two woredas were purposefully selected depending on the security and sheep production potential of the areas. From each woreda, two kebeles were selected purposively, and study animals were selected using simple random methods from healthy and purposively from diseased sheep (those showing signs of pasteurellosis) brought to the woreda’s veterinary clinic during the visit.

### Sample Size Determination

Since the prevalence of ovine pasteurellosis in the study areas was not known, the sample size was determined using the formula given by ^18^ based on a maximum expected prevalence of 50%. For this study, a level of significance of 95% is considered. Hence,

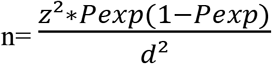Where,

n=number of sample size,

z= constant 1.96

P_exp_ expected prevalence of pneumonic pasteurellosis and,

d=desired absolute precision level.

Therefore, 384 local breeds of sheep were considered in the study. Sample size was allocated to each woreda according to the proportion of the number of sheep they possess. The total number of sheep from each study district was 72, 58, 70, 72, 50, and 62, from Horro Guduru Wollega (Jima-Geneti, Abe Dongoro), East Wollega (Wayu Tuqa, Diga), and West Shawa (Bako Tibe and Calliya) woreda, respectively.

### Specimen Collection and Transportation

Samples were obtained from a total of 384 sheep, comprising 220 healthy and 164 diseased individuals. Each sheep was assigned a unique identification number and properly restrained. The external nasal area of the selected sheep was sterilized using 70% alcohol. Bilateral nasal swabs were collected using sterile cotton swabs moistened with peptone water, which were subsequently placed in test tubes containing 3 mL of peptone water. The samples were then stored in an ice box and transported to the Animal Science Department Laboratory at Wallaga University, Shambu Campus, following the protocol described in a previous study ^26^.

### Isolation and Identification of bacterial isolates

#### Bacteriological Identification

The isolation and identification of *Pasteurella* were conducted at the Shambu Campus Animal Science Laboratory using methods recommended by Hardy Diagnostics, USA^8^. Nasal swabs were incubated in tryptose soy broth (HIMEDIA, India) at 37 °C for 24 hours, followed by streaking on blood agar (OXOID, UK) enriched with 5% defibrinated sheep blood and incubated aerobically at 37 °C for another 24 hours.

Culture-positive colonies were examined through Gram staining to observe staining characteristics and cellular morphology under a 100x light microscope. Gram-negative bacteria indicative of *Pasteurella* were subcultured onto blood agar and MacConkey agar (HIMEDIA, India). Colony characteristics, such as hemolysis type, morphology, color, shape, size, consistency, and lactose fermentation ability, were assessed on blood agar^7,8^.

Pure cultures were transferred to nutrient agar slants for biochemical tests, including catalase (Fisher Chemical, UK), indole (HIMEDIA, India), urease (HIMEDIA, India), triple sugar iron (TSI; BIOMAKTM, India), and MR-VP tests (HIMEDIA, India), following MacFadyen’s methods ^29^.

The isolates were identified based on their characteristics: *M. haemolytica* showed β-hemolysis on blood agar, pink pinpoints on MacConkey agar, was catalase-positive, and indole-negative. *B. trehalosi* displayed a narrow hemolytic zone on blood agar, no pink pinpoints on MacConkey agar, was catalase-negative, and indole-negative. *P. multocida* showed no hemolysis on blood agar, mucoid colonies, no growth on MacConkey agar, was catalase-positive, and indole-positive.

Finally, presumptive *Pasteurella* and *Mannheimia* colonies were preserved in nutrient broth with 98% glycerol and sent to the National Agricultural Biotechnology Research Center (Holeta, Ethiopia) for molecular identification.

#### Molecular Identification

The molecular detection of the isolated colonies of *M. haemolytica* and *P. multocida* was conducted at the National Agricultural Biotechnology Research Center, which is located in Holeta town, Ethiopia, by the following procedure.

The DNA of *M. haemolytica* and *P. multocida* was extracted using the boiling method. A single colony was collected with a loop and re-suspended in 100 μL of nuclease-free water by swirling the loop in an Eppendorf tube, followed by vortexing for 30 seconds. The tube was then placed in a thermal block preheated to 95–100°C and incubated for 10 minutes. Afterward, the tube was cooled at room temperature for 2 minutes and centrifuged at 12,000 rpm for 5 minutes using a mini centrifuge. Subsequently, 50 μL of the supernatant was carefully transferred to a fresh tube, avoiding the pellet. This supernatant served as the template DNA and was stored at −20°C for future use ^30,31^.

Conventional PCR was performed simultaneously amplifying the two virulence-associated genes of *M. haemolytica, i*.*e*., *PHSSA (P. haemolytica* serotype-specific antigen), a serotype-specific antigen of *M. haemolytica*, and Rpt2, a gene coding for methyl transferase. The primer design used was according to ^32,^ with some modifications and sizes of amplified products provided in Table 1.

**Table 1:**
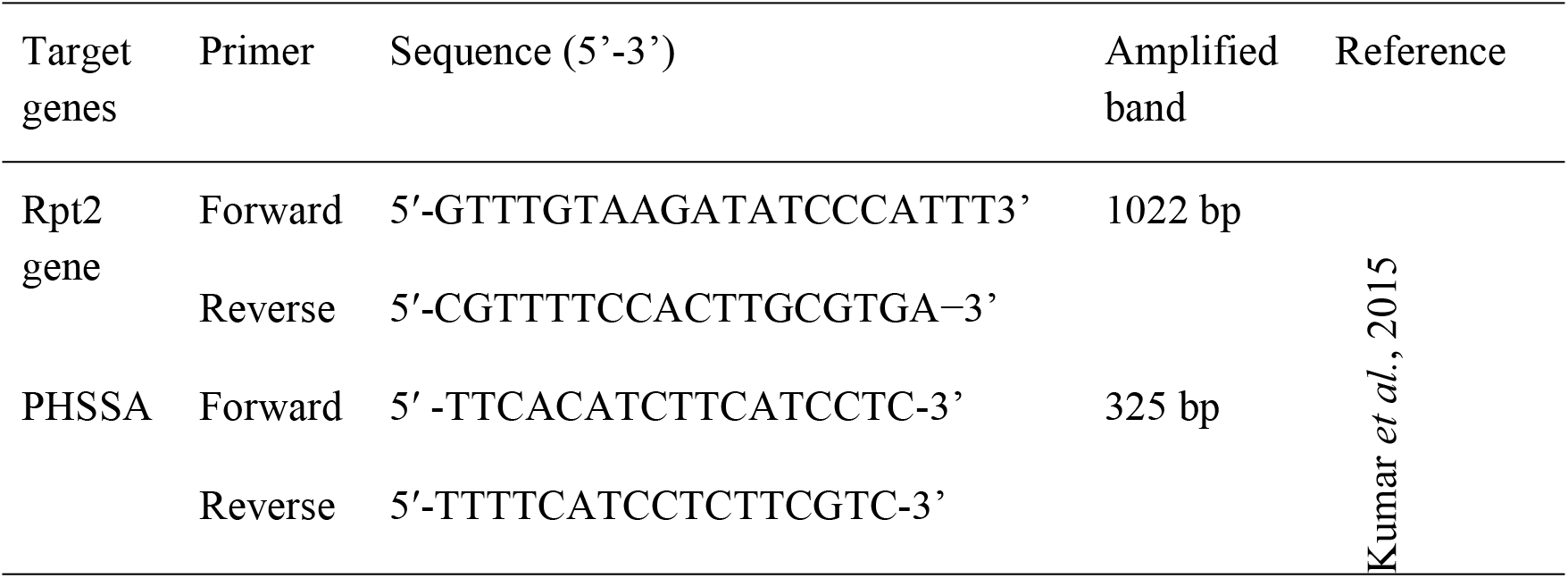
Primer (Forward and Reverse) used for *M. haemolytica*.

The PCR reaction mixture consisted of 10 μL of IQ Super-mix (Bio-Rad, USA), containing DNA polymerase, dNTPs, and buffer; 2 μL (5 pM/μL) of each primer pair; 3 μL of nuclease-free water; and 4 μL of DNA template, making a total reaction volume of 25 μL. The PCR conditions included an initial denaturation at 95 °C for 3 minutes, followed by 35 cycles of denaturation at 95 °C for 1 minute, annealing at 48 °C for 1 minute, and extension at 72 °C for 30 seconds. The final extension was carried out at 72 °C for 5 minutes.

A negative control (reaction without DNA template) and a positive control (reaction using the DNA template from the reference *M. haemolytica* isolate obtained from the National Veterinary Institute culture collection, MH-NVI) were included in the setup. Conventional PCR was performed to amplify the virulence-associated capsular gene (*CapA*) of *P. multocida*. The primers used were based on the design described in reference ^33^ with slight modifications. The sizes of the amplified products are summarized in Table 2 below.

**Table 2:**
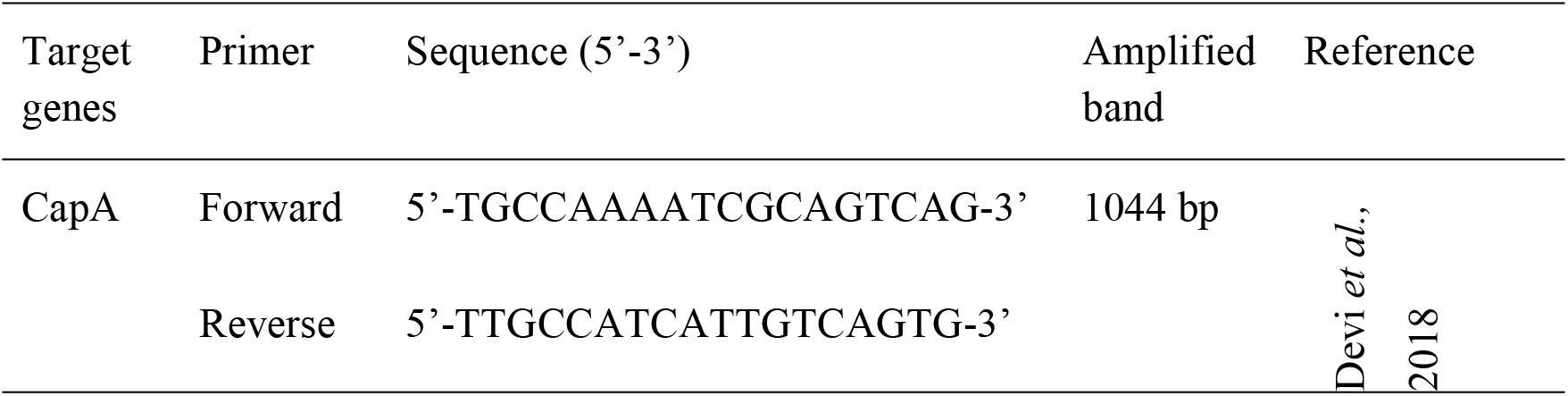
Primer (Forward and Reverse) used for *P. multocida*.

The PCR reaction was conducted in a total volume of 25 µL, comprising 5 µL of template DNA, 4.5 µL of nuclease-free water, 2 µL each of forward and reverse primers (5 pmol/µL; Eurofins MWG Operon, Germany), 5 µL of 10x DreamTaq™ Buffer containing 20 mM MgCl_2_, 5 µL of dNTP mix (2 mM each), and 1.5 µL of DreamTaq™ polymerase. The reaction conditions included an initial denaturation at 95 ºC for 5 minutes, followed by 35 cycles of denaturation at 95 ºC for 1 minute, annealing at 55 ºC for 1 minute, extension at 72 ºC for 30 seconds, and a final extension at 72 ºC for 7 minutes. A positive control DNA was obtained from the *P. multocida* vaccine strain, while a negative control, consisting of the reaction mixture without the DNA template, was included in the assay.

#### Agarose gel analysis of the polymerase chain reaction products

Agarose gel (2% w/v in 0.5 x Tris-borate EDTA buffer) stained with GelRed was employed for the analysis and detection of the PCR products. A 5 µl aliquot of each PCR product was loaded into separate wells of the prepared gel, along with 6x loading dye. A 1kb molecular marker was loaded in the first lane, and electrophoresis was conducted at 120V for 60 minutes using an electrophoresis apparatus (EC, 2060, USA). Subsequently, the resulting bands were captured using a gel documentation system and visualized under a UV transilluminator.

### Data Management and Analysis

Microsoft Excel was used to enter the field and lab data, and R software versio**n** 4.2.2 was used for analysis. Sums and frequency distributions were among the descriptive statistics that were calculated. Using logistic regression and chi-square (χ^2^) testing, associations between risk factors and the incidence of Pasteurella and Mannheimia in sheep were assessed. Both univariate and multivariable logistic regression models were used to determine odds ratios (OR). A stepwise logistic regression model with a significance level of P < 0.05 was used once the data had been examined for assumption fulfillment. The Hosmer–Lemeshow goodness-of-fit test was used to evaluate the model’s fitness; a good fit was indicated by a P > 0.05 value. A 95% confidence level was used for all analyses. ArcGIS 10.7 was used to map the study region.

## Results

The prevalence of *P. multocida, M. haemolytica*, and *B. trehalosi* was assessed in sheep categorized as either healthy or diseased. The results revealed a markedly higher prevalence of *P. multocida* among diseased sheep (18.9%) compared to healthy sheep (2.7%). The odds of isolating *P. multocida* from diseased animals were more than eight times higher than from healthy ones (OR = 8.31, 95% CI: 3.38–20.46, p < 0.0001), indicating a statistically significant association between *P. multocida* infection and diseased status (Table 3).

**Table 3.** Prevalence and association of *P. multocida, M. haemolytica*, and *B. trehalosi* with the health status of sheep.

| Pathogen | Status of Sheep | No. Examined | Positive n (%) | OR (95% CI) | p-value |
| --- | --- | --- | --- | --- | --- |
| *P. multocida* | Healthy | 220 | 6 (2.7%) | Ref | <0.01** |
|  | Diseased | 164 | 31 (18.9%) | 8.31 (3.38–20.46) |  |
| *M. haemolytica* | Healthy | 220 | 11 (5.0%) | Ref | 0.110 |
|  | Diseased | 164 | 15 (9.1%) | 1.91 (0.85–4.28) |  |
| *B. trehalosi* | Healthy | 220 | 12 (5.5%) | Ref | 0.410 |
|  | Diseased | 164 | 6 (3.7%) | 0.66 (0.24–1.79) |  |

For *M. haemolytica*, the prevalence was 9.1% in diseased sheep and 5.0% in healthy sheep. Although the odds ratio suggested nearly twice the risk in diseased animals (OR = 1.91, 95% CI: 0.85–4.28), the association was not statistically significant (p = 0.110).

In contrast, *B. trehalosi* was detected in 5.5% of healthy sheep and 3.7% of diseased sheep. The odds of isolation were lower in diseased animals (OR = 0.66, 95% CI: 0.24–1.79), but the association was not statistically significant (p = 0.410). These findings highlight that *P. multocida* is significantly associated with disease status in sheep, while *M. haemolytica* and *B. trehalosi* do not show a statistically meaningful relationship.

The prevalence of isolated species was compared across six districts. The overall prevalence was 21.1% (81/384). Among the districts, Bako Tibe had the highest prevalence at 26.0% (13/50), followed by Abe Dongoro with 22.4% (13/58). Diga and Wayu Tuqa recorded prevalences of 20.8% and 20.0%, respectively, while J/Geneti and Calliya both had the lowest prevalence at 19.4% (Table 4).

**Table 4.** Summary of the frequency of isolated species in different districts.

| Woreda | No. Examined | Positive (n) | Prevalence (%) | $\chi^2$ | df | p-value |
| --- | --- | --- | --- | --- | --- | --- |
| J/Geneti | 72 | 14 | 19.4% | 1.067 | 5 | 0.957 |
| Abe Dongoro | 58 | 13 | 22.4% |  |  |  |
| Wayu Tuqa | 70 | 14 | 20.0% |  |  |  |
| Diga | 72 | 15 | 20.8% |  |  |  |
| Bako Tibe | 50 | 13 | 26.0% |  |  |  |
| Calliya | 62 | 12 | 19.4% |  |  |  |
| **Total** | **384** | **81** | **21.1%** |  |  |  |

Although there were slight variations in prevalence among the districts, statistical analysis showed that the differences were not significant (χ^2^ = 1.067, df = 5, p = 0.957). This suggests that the distribution of the isolates was relatively uniform across the study areas, with no district showing a significantly higher or lower risk of isolation.

The overall prevalence of *Pasteurellaceae* in sheep across the study area was 21.1% (81/384). When stratified by geographic zone, the prevalence was slightly higher in West Shawa (22.3%) compared to H/G/Wallaga (20.8%) and East Wallaga (20.4%). However, no statistically significant differences were observed between zones, as indicated by the odds ratios and confidence intervals (H/G/Wallaga: OR = 0.98, 95% CI = 0.54–1.76, P = 0.944; West Shawa: OR = 0.89, 95% CI = 0.49–1.63, P = 0.713), suggesting that geographic location did not have a significant effect on *Pasteurellaceae* isolation (Table 5).

**Table 5.**
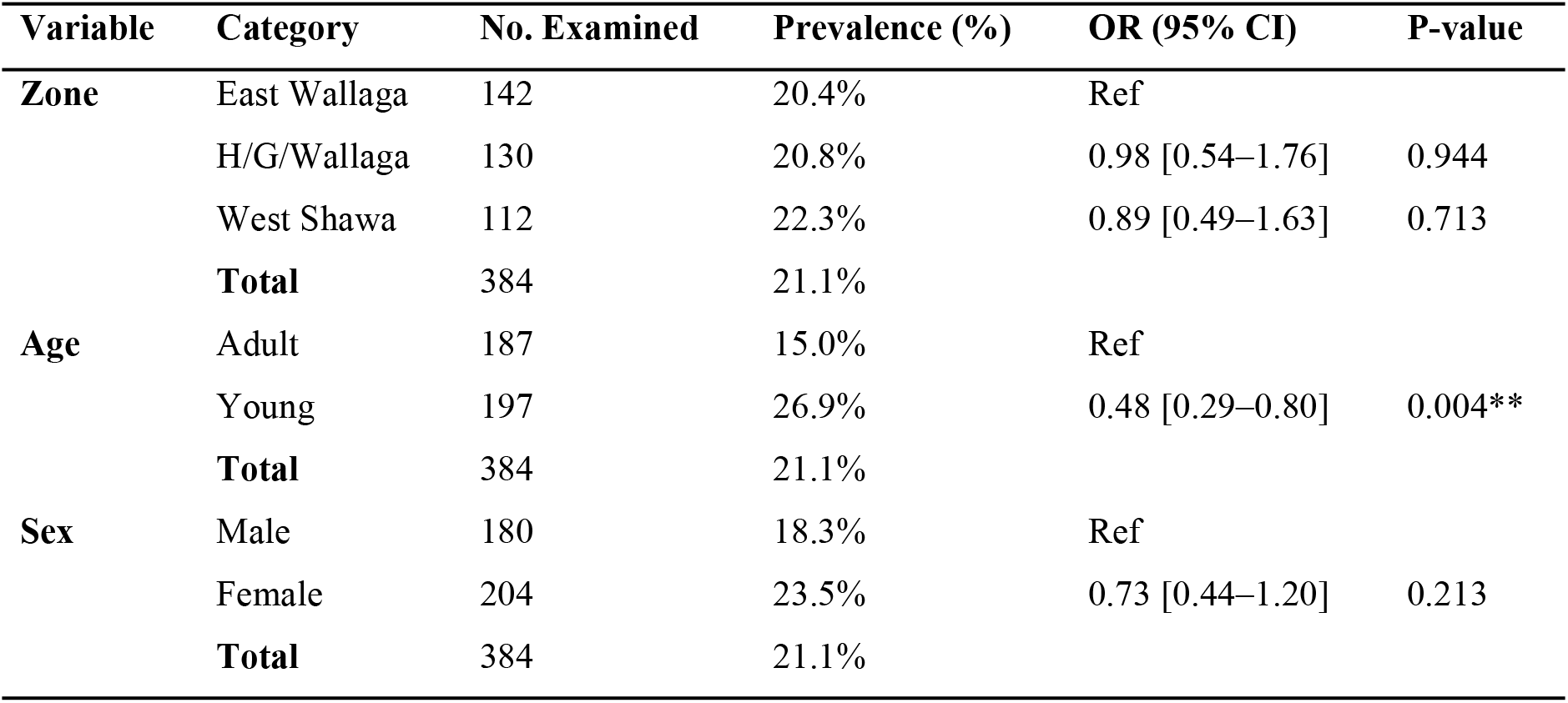
Prevalence of isolated Pasteurellaceae species by zone, age, and sex of sheep in Western Oromia, Ethiopia.

Stratification by age revealed a markedly higher prevalence among young sheep (26.9%) compared to adults (15.0%). The difference was statistically significant, with young animals being more than twice as likely to harbor *Pasteurellaceae* than adults (OR = 0.48, 95% CI = 0.29–0.80, P = 0.004), indicating that age is an important risk factor for infection.

Regarding sex, female sheep exhibited a higher prevalence (23.5%) than males (18.3%). However, the association was not statistically significant (OR = 0.73, 95% CI = 0.44–1.20, P = 0.213), suggesting that sex did not substantially influence the likelihood of *Pasteurellaceae* isolation in this population.

Overall, these findings indicate that age is a significant determinant of *Pasteurellaceae* prevalence in sheep, whereas geographic zone and sex appear to have a minimal effect in the study area.

Multi-logistic regression analysis revealed that age, sex, and health status were associated with the prevalence of *Pasteurellaceae* species (Table 6). Young animals were significantly more susceptible than adults (OR = 2.41; 95% CI: 1.42–4.18; p = 0.001), indicating higher vulnerability due to immature immune systems. Female animals exhibited a slightly higher risk compared to males (OR = 1.67; 95% CI: 0.99–2.86; p = 0.050), suggesting a possible sex-related predisposition. Notably, animals with pneumonic conditions had a markedly higher likelihood of infection than non-pneumonic animals (OR = 3.47; 95% CI: 2.06–5.97; p < 0.001), emphasizing the strong link between respiratory disease and pathogen prevalence. These results highlight the need to prioritize young and pneumonic animals in control and prevention strategies for *Pasteurellaceae* infections.

**Table 6.** Multi-logistic regression analysis of *Pasteurellaceae* species with different risk factors.

| Predictor | Category | Prevalence, n (%) | OR [95% CI] | P-value |
| --- | --- | --- | --- | --- |
| Age | Adult | 28 (7.3) | Ref | — |
|  | Young | 53 (13.8) | 2.41 [1.42–4.18] | 0.001** |
| Sex | Male | 38 (8.6) | Ref | — |
|  | Female | 48 (12.5) | 1.67 [0.99–2.86] | 0.050 |
| Health Status | Non-pneumonic | 29 (7.55) | Ref | — |
|  | Pneumonic | 52 (13.55) | 3.47 [2.06–5.97] | <0.001** |

Out of 26 isolates identified as *Mannheimia haemolytica* by bacteriological and biochemical tests, seven representative samples were subjected to PCR for the detection of the PHSSA and Rpt2 genes. Among these, five samples (71.4%; 95% CI: 0.36–0.92) were found to be positive, while two were negative.

Similarly, out of 37 isolates identified as *Pasteurella multocida*, 16 were examined for the presence of the CapA gene, of which five (31.3%; 95% CI: 0.14–0.56) tested positive and 11 were negative.

Figure 3 shows the results of PCR amplification of a *P. multocida* product (1044 bp). Lanes 9, 10, 11, 13, and 14 are *P. multocida*.

**Figure 1.**
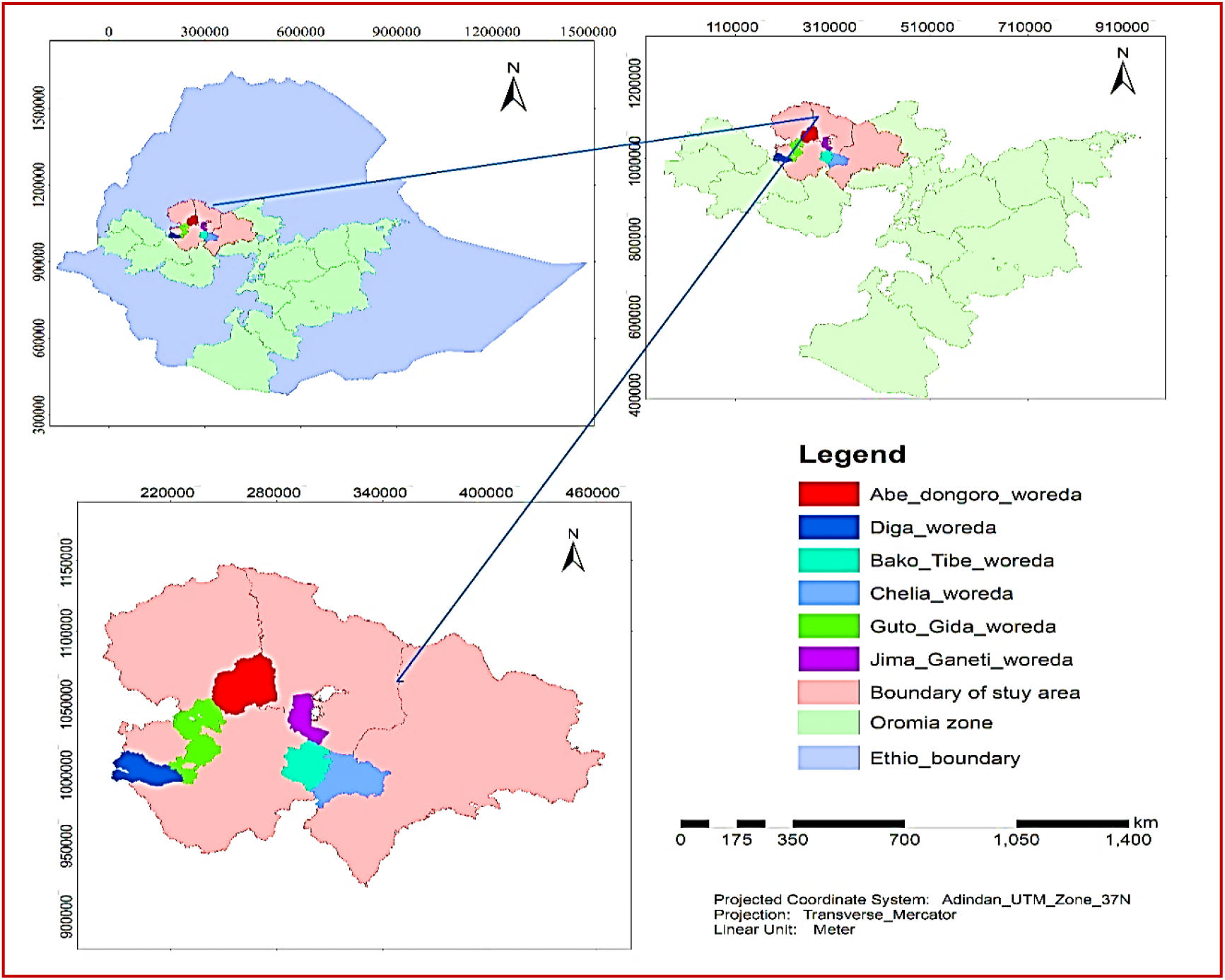
Map of the study area (Source: ArcGIS version 10.7)

**Figure 2.**
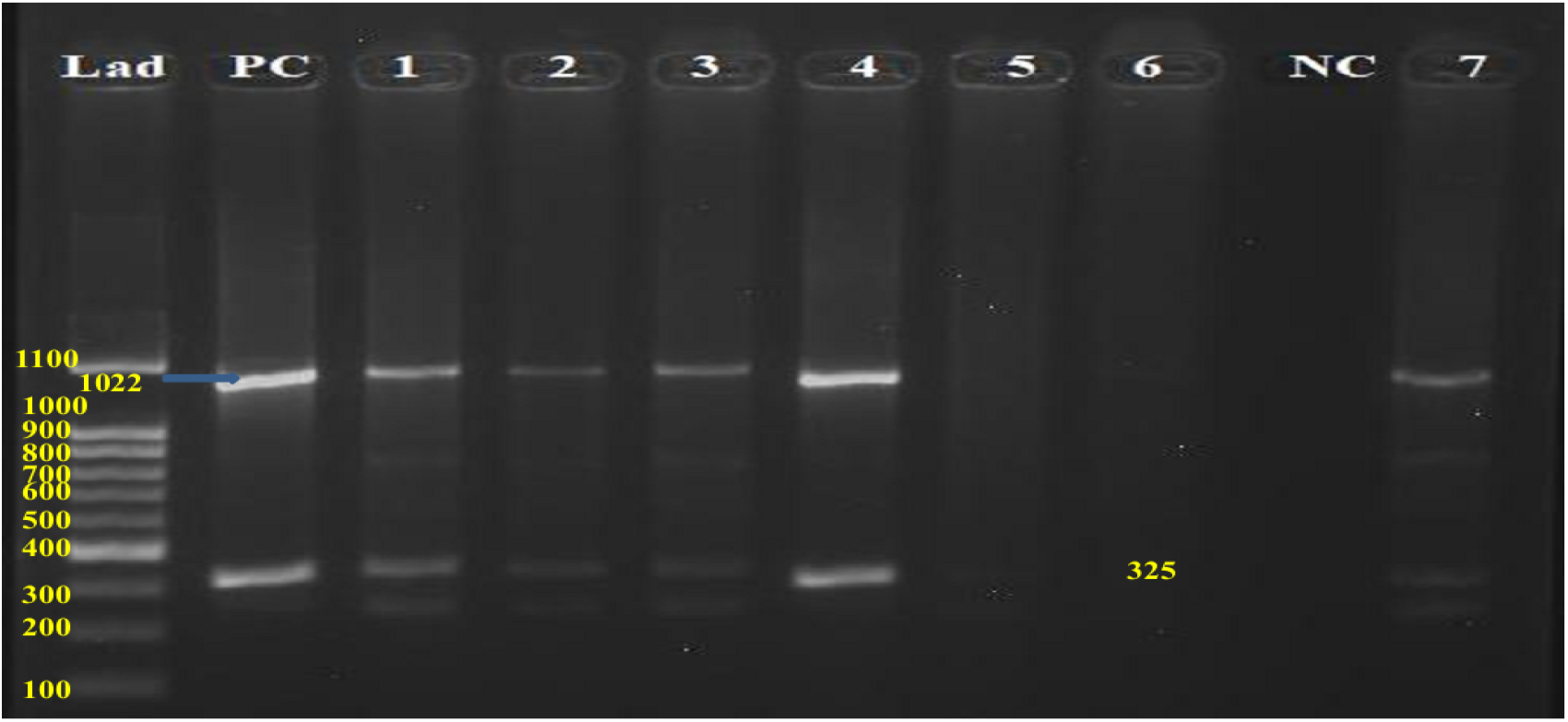
Gel electrophoresis of amplified product of PHSSA and Rpt_2_ Hint: Lad=Ladder, PC=Positive control, 1-7=Test isolates, NC=Negative control

**Figure 3.**
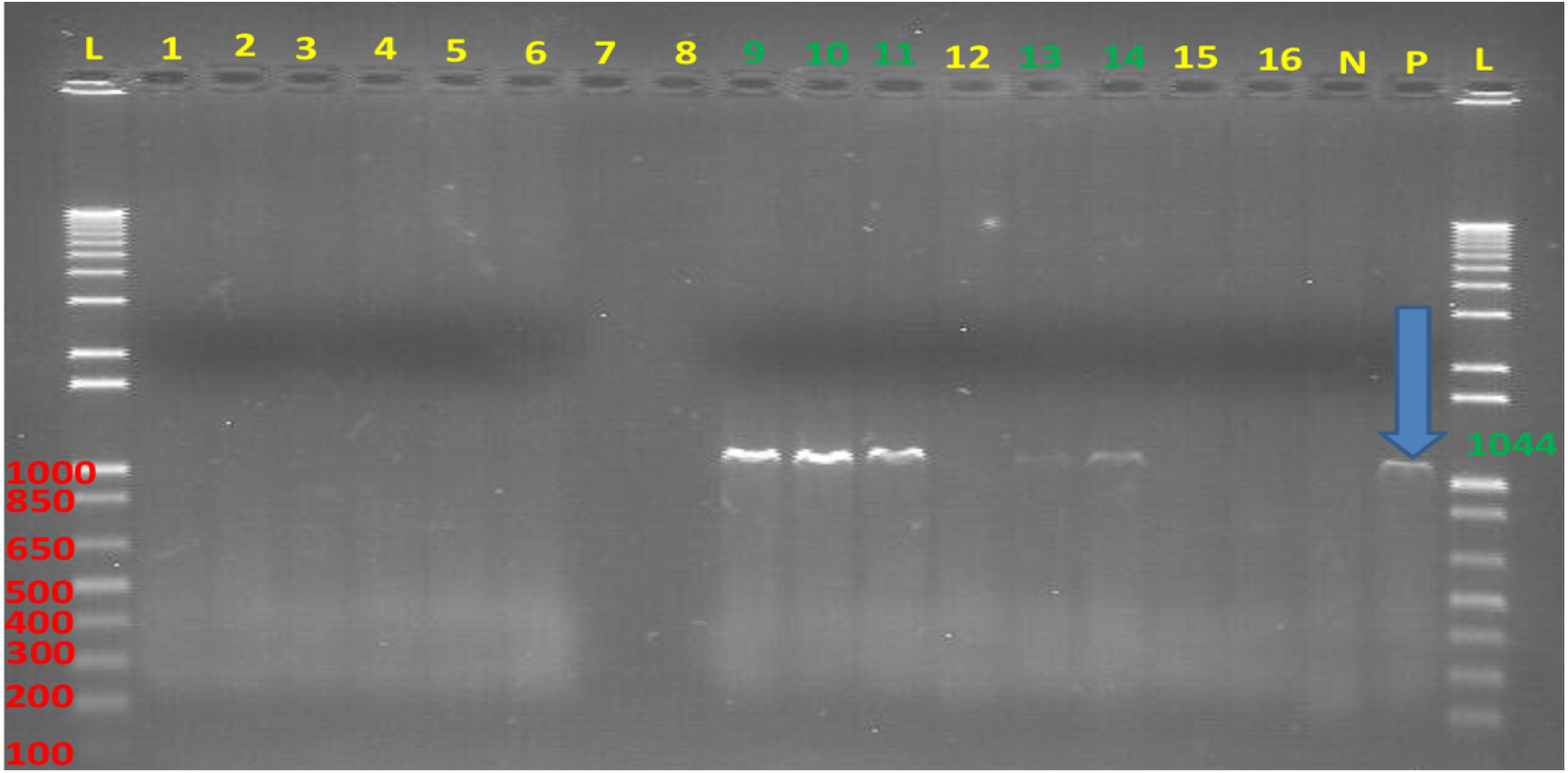
Gel electrophoresis of amplified product of CapA gene *Hint*:-L= Ladder; N= negative control; P= positive control; 1-16= isolates of bacteria.

In contrast, *Bibersteinia trehalosi* (n = 18) was not included in molecular testing due to the unavailability of specific primers (Table 7).

**Table 7.** Table: Molecular Detection of Pasteurellaceae Species.

| Species | Total Isolates<br>(Biochemical) | Isolates<br>Subjected to<br>PCR | Positive<br>(%) | 95% CI<br>(Proportion) | Chi-square | P-value |
| --- | --- | --- | --- | --- | --- | --- |
| *M. haemolytica* | 26 | 7 | 71.4%<br>(5/7) | 0.36 – 0.92 |  |  |
| *P. multocida* | 37 | 16 | 31.3%<br>(5/16) | 0.14 – 0.56 | 1.77 | 0.183 |
| *B. trehalosi* | 18 | 0 | – | – | – | – |

When the positivity rates of *M. haemolytica* and *P. multocida* were compared, a higher proportion of positive samples was observed in *M. haemolytica* (71.4%) compared to *P. multocida* (31.3%). However, this difference was not statistically significant (χ^2^ = 1.77; p = 0.183), indicating that the variation in detection rates between the two species could be due to chance rather than a true biological difference.

## Discussion

The overall prevalence of Bibersteinia trehalosi, Mannheimia hemolytica, and Pasteurella multocida in ovine pasteurellosis was 21.1%, according to the current study. Previous reports from other regions of Ethiopia are consistent with this finding. In sheep from Lume District, East Shewa Zone, a similar incidence of 24.1% was observed by Haji et al. (2016). Prevalence rates in the districts of Asella, Holota, and Sheno were also reported by Wirtu et al. (2022) to be 25%, 26.5%, and 23.5%, respectively. Marru et al. (2013) reported similar results in Haramaya District, Eastern Hararghe, where they reported a 25% prevalence of pneumonic pasteurellosis, which was dominated by M. haemolytica. Furthermore, Asefa (2012) reported that sheep from particular districts in the Arsi Zone, Oromia Region, had notable isolation rates of M. haemolytica (59.6%), P. multocida (30.3%), and B. trehalosi (10.1%). All of these results show that ovine pasteurellosis is still a significant respiratory condition that is common in Ethiopia ^34, 35, 36, 37^.

The current prevalence (21.1%) is higher than what has been observed in a number of other nations when compared globally. Sahay et al. (2020) reported a prevalence of M. haemolytica and *P. multocida* together of around 14.7% of sampled sheep (7.2% *M. haemolytica*, 7.4% *P. multocida*) in Karnataka, southern India ^38^. This is significantly lower than ours. A study conducted in the Kars Province of Turkey (Dağ et al., 2018) found that *P. multocida* was present in 3% of pneumonic lung samples and *M. haemolytica* in 19% ^39^. *M. haemolytica* accounted for approximately 76.8% and *P. multocida* for approximately 23.2% of the 42.7% overall prevalence of pasteurellosis in small ruminants, according to a recent study conducted in the Haramaya District in eastern Ethiopia (2024) ^40^.

A slightly significant sex difference was found: females had a higher prevalence, which may have been brought on by increased physiological stressors during pregnancy, parturition, and lactation. Similar trends have been documented in other places. In one Ethiopian study, for example, Mannheimia haemolytica was discovered in 14.86% of females and 9.46% of males ^41^. Furthermore, studies conducted in Ethiopia showed that testing positive was more common in girls (36.42%) than in males (37.1%) ^42^.

Significant correlations were found between age and infection: the prevalence was higher in lambs and young sheep (13.8%) than in adults (7.3%), which is in line with research showing that younger animals are more susceptible ^43^. Disease status also affected the frequency of isolations: isolates were more common in sheep with clinical disease (13.5%) compared to sheep that appeared to be healthy (7.5%). This suggests that these bacteria are frequently recovered from pneumonic animals and can function as primary or secondary pathogens ^43, 40^. Ultimately, our results highlight the probable role of *B. trehalosi, M. haemolytica*, and *P. multocida* in pneumonic pasteurellosis and their potential to affect ruminant productivity nationally.

Sheep health status and the prevalence of Pasteurella and Mannheimia species were shown to be significantly correlated. The prevalence of diseased sheep was higher (13.5%) than that of sheep that appeared to be in good health (7.5%). This is in line with results from other research, including one carried out in Ethiopia, which found that *M. haemolytica* and *B. trehalosi* were more common in pneumonic sheep than in healthy ones ^44^. In a similar vein, *M. haemolytica* was found in 46.8% of the sheep studied in a California research, suggesting that it may have a role in respiratory infections ^45^. These results indicate the possible role of Pasteurella and Mannheimia species as main or secondary pathogens in respiratory illnesses, despite the fact that they are common commensals in the upper respiratory tract and that they are more frequently isolated in pneumonic animals.

## Conclusion And Recommendations

Pneumonic pasteurellosis is the most economically important infectious disease of sheep throughout the world. The present study report was an attempt to isolate, identify, and molecularly detect *P. multocida, M. haemolytica*, and *B. trehalosi from both apparently healthy and diseased sheep in selected areas of western Oromia*. The study revealed that *P. multocida, M. haemolytica*, and *B. trehalosi were isolated from sheep in the study areas. In the present study, P. multocida is the most prevalent isolated species, followed by M. haemolytica and B. trehalosi*. The serotypes confirmed by molecular methods of *P. multocida and M. haemolytica* are those serotypes that cause pneumonia in ovine (*P. multocida serotype A* and *M. haemolytica*). Furthermore, understanding the disease’s prevalence at the species level aids in the development of the appropriate intervention strategy, whether it be a vaccination, antibiotics, or another alternative technique.

This study’s conclusions lead to a number of recommendations. To precisely identify and validate the isolates’ species or serotypes, molecular techniques should be used in future research. Moreover, as certain significant locations were left out of the current study, more investigation is required to examine additional places and to molecularly and phylogenetically describe the etiological agents linked to ovine pasteurellosis. Furthermore, as Pasteurella and Mannheimia species were isolated from both clinically ill and seemingly healthy sheep, their precise function in the development of pneumonia merits more research. A better grasp of their pathogenic relevance and the development of efficient control measures might result from such investigations.

## Notes

### Competing Interest Statement

The authors have declared no competing interest.

## References

1. Zeleke, B. (2017). Status and growth trend of draught animals’ population in Ethiopia. Journal of Dairy, Veterinary & Animal Research, 6 (1), 238–241.

2. Abebe, G. (2022). Evaluation of Microbial Safety and Physicochemical Quality of Carcas from Borena and Central Highland Of North Shewa Goats (Doctoral Dissertation). Retrieved from http://ir.bdu.edu.et/handle/123456789/14720.

3. Asresie, A., Zemedu, L., and Adigrat, E. (2015). The contribution of livestock sector in Ethiopian economy. A Review Advances in Life Science and Technology, 29.

4. Abera, D., and Mossie, T. (2023). A review on pneumonic pasteurellosis in small ruminants. Journal of Applied Animal Research, 51(1), 1–10.

5. Jilo, K., Belachew, T., Birhanu, W., Habte, D., Yadete, W., and Giro, A. (2020). Pasteurellosis Status in Ethiopia: A Comprehensive Review. J Trop Dis 8:351.

6. Donachie W. (2000). Pasteurellosis; Diseases of sheep. Blackwell Scientific Oxford, pp: 191–197.

7. Confer, A.W. (2009). “Update on Bacterial Pathogenesis in BRD.” Anim. Health Res. Rev.10 (2): 145–8.

8. Afata AW. (2018). Isolation and identification of Mannheimia haemolytica, Bibersteinia trehalosi and Pasteurella multocida from cattle and sheep from selected areas of Ethiopia, MSc Thesis, Addis Ababa University, Ethiopia.

9. Akane, A. E., Alemu, G., Tesfaye, K., Ali, D. A., Abayneh, T., Kenubih, A. and Ibrahim, S.M. (2022). Isolation and Molecular Detection of Pasteurellosis from Pneumonic Sheep in Selected Areas of Amhara Region, Ethiopia: An Implication for Designing Effective Ovine Pasteurellosis Vaccine. Veterinary Medicine: Research and Reports, 13, 75.

10. Ievy, S., Khan, M., Islam, M., and Rahman, M. (2013). Isolation and identification of Pasteurella multocida from chicken for the preparation of oil adjuvanted vaccine. Microbes and Health, 2 (1): 1–4.

11. Singh, F., G.G. Sonawane and R.K. Meena. (2018). Molecular detection of virulent M.haemolytica and P.multocidain lung tissues of pneumonic sheep from semiarid tropics, Rajasthan, India. Turkish Journal of Veterinary and Animal Sciences, 42 (6): 556–561.

12. Jesse, F. F. A., Boorei, M. A., Chung, E. L. T., Wan-Nor, F., Lila, M. M., Norsidin, M. J. and Paul, B. T. (2020). A review on the potential effects of Mannheimia haemolytica and its immunogens on the female reproductive physiology and performance of small ruminants. Journal of Animal Health and Production.

13. Karimkhani, H., Zahraie, S. T., Sadeghi, Z. M., Karimkhani, M., and Lameyi, R. (2011). Isolation of pasteurella multocida from cows and buffaloes in Urmia’s slaughter house.

14. Haji, S. (2015). Pasteurollosis in sheep and its drug susceptibility pattern in Mojo district, East Shoa Zone (Doctoral dissertation, Addis Ababa University).

15. CSA. (2021). Agricultural Sample Survey 2020/21 (2013 E.C). Volume II report on livestock and livestock characteristics (private peasant holdings). Central Statistical Agency (CSA): Addis Ababa, Ethiopia.

16. Olana, B. T. (2006). People and dams: environmental and socio-economic changes induced by a reservoir in Fincha’a watershed, western Ethiopia. Wageningen University and Research.

17. Amante, M., Hunde, A., Endebu, B., Hirpa, E., and Mamo, B., (2014). Health and welfare assessment of working equine in and around Nekemte Town, East Wollega Zone, Ethiopia. American-Eurasian journal of scientific research, 9 (6), 163–174.

18. Tamiru, F., M., Hailemariam, and W., Terfa. (2013). Preliminary study on prevalence of bovine tuberculosis in cattle owned by tuberculosis positive and negative farmers and assessment of zoonotic awareness in Ambo and Toke Kutaye districts, Ethiopia. Journal of Veterinary Medicine and Animal Health, 5 (10), 288–295.

19. Tesfa, M., Sualeh, A., and Mekonen, N. (2021). Assessment of the Effectiveness of Coffee De-mucilager and Driers for Physical and Sensorial Coffee Quality. World, 5 (2), 33–36.

20. Megersa, M., and Tamrat, N. (2022). Medicinal Plants Used to Treat Human and Livestock Ailments in Basona Werana District, North Shewa Zone, Amhara Region, Ethiopia. EvidenceBased Complementary and Alternative Medicine, 2022.

21. Duressa, D., Kenea, D., Keba, W., Desta, Z., Berki, G., Leta, G., and Tolera, A. (2014). Assessment of livestock production system and feed resources availability in three villages of Diga district Ethiopia.

22. Abera, Z., Fekadu, M., Kabeta, T., Kebede, G., and Mersha, T. (2014). Prevalence of bovine trypanosomosis in bako tibe district of west shoa and gobu seyo districts of west wollega Zone, Ethiopia. Eur J Biol Sci, 6(3), 71–80.

23. Degefa, K., Biru, G., and Abebe, G. (2020). Characterization and analysis of farming system of Cheliya and Ilu Gelan districts of West Shewa Zone, Ethiopia. Journal of Development and Agricultural Economics, 12 (3), 154–167.

24. Gatenby, M.R. (1991): sheep. Coste, R. and Smith, J.A. (eds), the tropical agriculturalist, Macmillan (London) and CTA (Wageningen), Pp. 6–11.

25. Thrusfield, M. (2018). Veterinary epidemiology. John Wiley & Sons.

26. Marru, H. D., Anijajo, T. T., and Hassen, A. A. (2013). A study on Ovine pneumonic pasteurellosis: Isolation and Identification of Pasteurellae and their antibiogram susceptibility pattern in Haramaya District, Eastern Hararghe, Ethiopia. BMC Veterinary Research, 9 (1), 1–8.

27. Tabatabaei, M., and Abdollahi, F. (2018). Isolation and identification of Mannheimia haemolytica by culture and polymerase chain reaction from sheep’s pulmonary samples in Shiraz, Iran. Veterinary World, 11 (5), 636.

28. Mahon, C. R. (2022). Use of colony morphology for the presumptive identification of microorganisms. Textbook of Diagnostic Microbiology-E-Book, 169.

29. MacFadinn, J.F. (2000). Biochemical tests for identification of medical bacteria. 3rd edition. New York: Williams and Wilkins Lippincott. ISBN::0683, 05318-05183.

30. Dimitrakopoulou, M. E., Stavrou, V., Kotsalou, C., and Vantarakis, A. (2020). Boiling extraction method vs commercial kits for bacterial DNA isolation from food samples. Journal of Food Science and Nutrition Research, 3 (4), 311–319.

31. Ewers, C., Lubke, A., Becher, and Wieler, L.H. (2005). Mannheimia haemolytica and the pathogenesis of pneumonic pasteurellosis. Berliner and Munchener Veterinary Lice Wochenschrift, 117: 97–115.

32. Kumar, J., Dixit, S.K., Kumar, R. (2015). Rapid detection of M. haemolytica in lung tissues of sheep and from bacterial culture. Vet. World. 1073–1077.

33. Devi, L.B., Bora, D. P., Das, S. K., Sharma, R. K., Mukherjee, S., and Hazarika, R. A. (2018). Virulence gene profiling of porcine Pasteurella multocida isolates of Assam. Veterinary world: 11 (3), 348.

34. Haji, H., & Abunna, F. (2016). Epidemiology of ovine pasteurollosis in Lume district, East shewa zone of Oromiya region, Ethiopia. Int Know Share Plat, 6, 12–20.

35. Wirtu A, Kumsa B, Zerabruk E, Albene Y, Tadesse F, et al. (2022) Bacteriological and Molecular Identification of Mannheimia haemolytica, Pasteurella multocida and Bibersteinia trehalosi from Cattle and Sheep from Selected Areas of Ethiopia. J Vet Med Res 9(3): 1234.

36. Marru, H.D., Anijajo, T.T. & Hassen, A.A. A study on Ovine pneumonic pasteurellosis: Isolation and Identification of Pasteurellae and their antibiogram susceptibility pattern in Haramaya District, Eastern Hararghe, Ethiopia. BMC Vet Res 9, 239 (2013). 10.1186/1746-6148-9-239

37. Assefa, B. (2012). Epidemiology and Drug Resistance of Ovine Pasteurellosis In Selected Districts Of Arsi Zone Of Oromia Regional State, Ethiopia (Doctoral dissertation, MSc Thesis, Addis Ababa University, Debre Zeit, Ethiopia).

38. Sahay S, Natesan K, Prajapati A, Kalleshmurthy T, Shome BR, Rahman H, Shome R (2020) Prevalence and antibiotic susceptibility of Mannheimia haemolytica and Pasteurella multocida isolated from ovine respiratory infection: A study from Karnataka, Southern India, Veterinary World, 13(9): 1947–1954.

39. Dag, S., Gurbuz, A., Ozen, H., Buyuk, F., Celebi, O., Karaman, M., … ÇİTİL, M. (2018). Immunohistochemical and Molecular Detection of Mannheimia spp. and Pasteurella spp. in Sheep with Pneumonia in Kars Province - Turkey. KAFKAS UNIVERSITESI VETERINER FAKULTESI DERGISI, vol.24, no.2, 281–288.

40. Abdulkadir, M., Nigussie, T., & Kebede, I. A. (2024). Isolation and identification of Pasteurella multocida and Mannheimia haemolytica from pneumonic small ruminants and their antibiotic susceptibility in Haramaya District, Eastern Ethiopia. The Scientific World Journal, 2024(1), 5605552.

41. Abate FM, Fentie Kassa T. Isolation and identification of Mannheimia haemolytica and Pasteurella multocida from symptomatic and asymptomatic sheep and their antibiotic susceptibility patterns in three selected districts of north Gondar zone, Gondar Ethiopia. Vet Med Sci. 2023 Jul;9(4):1803–1811. doi: 10.1002/vms3.1166. Epub 2023 May 17. PMID: 37197762; PMCID: PMC10357255.

42. Alemneh, T., & Tewodros, A. (2016). Sheep and goats pasteurellosis: Isolation, identification, biochemical characterization and prevalence determination in Fogera Woreda, Ethiopia. Journal of Cell and Animal Biology, 10(4), 22–29.

43. Berhe, K., Weldeselassie, G., Bettridge, J., Christley, R. M., & Abdi, R. D. (2017). Small ruminant pasteurellosis in Tigray region, Ethiopia: marked serotype diversity may affect vaccine efficacy. Epidemiology & Infection, 145(7), 1326–1338.

44. Girma, S., Getachew, L., Beyene, A. et al. Identification of serotypes of Mannheimia haemolytica and Pasteurella multocida from pneumonic cases of sheep and goats and their antimicrobial sensitivity profiles in Borana and Arsi zones, Ethiopia. Sci Rep 13, 9008 (2023). 10.1038/s41598-023-36026-2

45. Jackson, W., Tucker, J., Fritz, H., Bross, C., Adams, J., Silva, M., Lorenz, C., & Marshall, E. (2024). Antimicrobial susceptibility profiles among commensal Mannheimia haemolytica and Pasteurella multocida isolated from apparently healthy sheep processed in California: Results from a cross-sectional pilot study. Preventive veterinary medicine, 233, 106360. 10.1016/j.prevetmed.2024.106360

